# Zika virus infection during pregnancy and induced brain pathology in beclin1-deficient mouse model

**DOI:** 10.1101/843813

**Authors:** Mohan Kumar Muthu Karuppan, Chet Raj Ojha, Myosotys Rodriguez, Jessica Lapierre, M. Javad Aman, Fatah Kashanchi, Michal Toborek, Madhavan Nair, Nazira El-Hage

**Author notes:** Correspondence: Dr. Nazira El-Hage, Department of Immunology and Nanomedicine, Florida International University, Herbert Wertheim College of Medicine, Miami, FL 33199, USA.

## Abstract

We investigated the role of the autophagy protein, Beclin1, in the replication and disease of Zika virus (ZIKV) in pregnant dams and their offspring using Beclin1-deficient (Atg6^+/−^) and wild-type (Atg6^+/+^) mouse model infected with the Honduran (R103451), Puerto Rican (PRVABC59), and the Uganda (MR766) strains of ZIKV. Pregnant dams infected subcutaneously at embryonic stage (E)9 showed viral RNA in serum harvested at E13 and in various organs removed postmortem at E17. Subcutaneous infections with ZIKV also showed the vertical transmission of ZIKV from the placenta to embryos removed postmortem at E17. From the three isolates, R103451-infected Atg6^+/−^ dams had the lowest mortality rate while 30 % of their offspring containing the hemizygous beclin1 allele (Atg6^+/−^) were smaller in size and had smaller and underdeveloped brain. Growth impairment in the pups became noticeable after two weeks post-birth. After 21-days, pups were sacrificed and brain tissues removed postmortem showed expression of the envelope (E) and the non-structural (NS)-1 proteins, along with signs of neuronal injury, despite an absence in viral RNA detection. A significant decrease in the mRNA expression levels of the insulin-like growth factor-1 (IGF-1) by 8-fold and a decrease in the mRNA expression levels of several microcephaly related genes along with an increase in the secretion of several inflammatory molecules may have contributed to the observed phenotype. Since autophagy regulates cytokines and chemokines production, a dysregulation in this pathway may have further exacerbated the pathology of ZIKV.

**IMPORTANCE:** Pups delivered from ZIKV-infected dams showed significant growth impairments in the body and the brain. We believe that the reduction in insulin growth factor together with the increase secretion of inflammatory molecules may have triggered neuronal injury and the downregulation of the microcephalic genes, while reduced expression of the autophagy protein, Beclin1 further exacerbated the pathology. Although the mechanism is still unknown, the autophagy pathway seems to play a key role in ZIKV pathology. It is therefore of great significance to study the role of autophagy during viral infection with the goal to identify potential targets for anti-ZIKV therapeutic intervention.

## INTRODUCTION

Zika virus (ZIKV) is a neurotropic flavivirus primarily transmitted by the Aedes mosquito (1–3). Individuals infected with the virus typically develop mild symptoms, although *in utero*, ZIKV exposure can cause congenital malformations including microcephaly (4, 5), and or other overt congenital abnormalities including fetal death, placental insufficiency, fetal growth restriction and central nervous system injury (6). Microcephaly is a neurodevelopmental disorder, characterized by a reduced head size when compared to babies of the same sex and age. The significant reduction in brain size, accompanied by intellectual disability, is believed to be caused by impaired cell proliferation and the death of cortical progenitor cells and their neuronal progeny (7). Significant downregulations of microcephaly-associated genes were detected in ZIKV-related studies (8–10), suggesting a direct mechanistic link of ZIKV infection to microcephaly. Twelve of the microcephalin (MCPH) loci (MCPH1-MCPH12) have been mapped (11) with many of these genes encoding for proteins localized at the centrosome or associated with centrosomal-related activities which play an important role in cell cycle progression, cell division and formation of the mitotic spindle (12).

Recently we have shown that ZIKV can modulate the autophagy pathway in glia (astrocytes and microglia) with silencing of the autophagy gene, *beclin1*, leading to increased inflammation in ZIKV-infected glia (13). Beclin1 is a component of the phosphatidylinositol 3-kinase nucleation complex which regulates the initiation stages of the autophagy pathway (14). Here we confirm our previous results using a translatable animal model. The Atg6^+/−^ mice expresses about 60% less Beclin1 protein (15) and serves as a valuable tool to analyze the function of Beclin1. We report for the first time that three different phylogenetic strains including the Honduran-R103451, Puerto Rican-PRVABC59 and the Uganda-MR766 strain of ZIKV infect the Atg6^+/−^ and Atg6^+/+^ pregnant dams. High mortality was detected in Atg6^+/+^ dams infected with MR766 when compared to dams infected with the Honduran-R103451 and the Puerto Rican-PRVABC59 strains, while R103451-infected animals showed the highest survival rate. Impaired growth in body and brain sizes were visible in 30 % of offspring born to R103451-infected Atg6 ^+/−^ dams, with no evidence of viral RNA in serum or brain removed at day-21 postnatal, albeit viral proteins were expressed in brain tissues. Significant reduction in IGF-1 along with signs of neuronal injury were detected in the brain of these pups. Furthermore, a significant downregulation in the expression levels of several microcephalic genes were evident, although decreased expression levels were also detected in brains of pups exposed to MR766 and PRVABC59 *in utero*.

## MATERIALS AND METHODS

### Viral propagation

Vero cells (CRL-1586), mosquito cell line C6/36 (CRL-1660), Honduran-R103451 (VR-1848) ZIKV, Puerto Rican-PRVABC59 (VR-1843) ZIKV and the Uganda-MR766 (VR84) ZIKV were procured from American Type Culture Collection (ATCC, Manassas, VA, USA). Vero cells or C6/36 were infected at multiplicity of infection (MOI) of 0.01 for propagation as described previously (13).

### Ethics statement

Animal work was conducted in accordance with the guidelines of the National Institutes of Health Guide for the Care and Use of Laboratory Animals. Animal experiments and associated protocols were reviewed and approved by the Florida International University Institutional Animal Care and Use Committee (IACUC).

### Animal model and timed pregnancy

*Atg6*^+*/*−^ (stock # 018429) mice and *Atg6*^+*/*+^ (stock # 000664) wild-type were procured from The Jackson Laboratory (Bar Harbor, ME, USA) and bred in the animal facility at Florida International University. For the timed-pregnancy studies, male and female mice aged 8 to 15 weeks were kept in isolation for at least 24 hours prior to mating. Males and females were placed together in the early evening and monitored periodically for up to 48 hours until the detection of a vaginal plug. The day plug was observed was considered embryonic or gestational day zero (E0). Pregnant dams received anti-interferon receptor 1 (anti-IFNAR1) monoclonal antibody (MAR-5A3, Leinco Technologies, MO, USA) at 2mg/animal via intraperitoneal (ip) route at gestational day 8 followed by subcutaneous (sc) infection with individual strain of ZIKV at 10^3^ plaque forming unit (PFU) in 50µL of PBS or mock (PBS) injection at gestational day of E9. Booster dose of anti-IFNAR1 at 0.5mg/animal dose was administered by ip at 2- and 4-days post-infection (dpi). At E13 (four days after ZIKV challenge), maternal blood was collected, and serum was prepared after coagulation and centrifugation. At E17, organs from dams (brain, liver, heart and spleen) placenta and fetuses were recovered. Organs were weighed and homogenized using a bead-beater apparatus (MagNA Lyser, Roche, Indianapolis). At E20-E21, pups delivered were monitored for growth and weight changes for up to 3 weeks of age.

### Real Time PCR

Viral RNA from serum and cellular RNA from tissues collected at various time-points from both ZIKV-infected and uninfected animals were extracted using QIAamp Viral RNA mini kit and RNeasy mini kit respectively (Qiagen, Valencia, CA, USA). The extracted RNA was amplified by iTaq universal SYBR Green one-step PCR kit (Bio-Rad, Hercules, CA, USA) and 10μM primers (Sigma-Aldrich, MO, USA). Expression of microcephalin-1 (MCPH1), WD repeat containing protein 62 (WDR62), cancer susceptibility candidate 5 (CASC5), and the abnormal spindle like primary microcephaly (ASPM) were measured using 500 ng of RNA extracted from pup brains. A standard curve was prepared from a 10-fold dilution of previously quantified ZIKV stock solution with known titer and viral titer expressed as Viral RNA (using standard curve). RT^2^ Profiler™ PCR Array Mouse Autophagy was purchased from Qiagen (Catalog # PAMM-084Z). RNA extracted from the brain of pups born to infected dams were analyzed for the expression of autophagy-related genes following the manufacturer’s instruction and as previously described (13).

### Primer sequences

#### ZIKV

Forward: 5’-CCGCTGCCCAACACAAG-3’

Reverse: 5’-CCACTAACGTTCTTTTGCAGACAT-3’

#### MCPH1

Forward 5′-AAGAAGAAAAGCCAACGAGAACA-3′

Reverse 5′-CTCGGGTGCGAATGAAAAGC-3’

#### ASPM

Forward 5′-CCGTACAGCTTGCTCCTTGT-3′

Reverse 5’-GGCGTTGTCCAATATCTTTCCA-3’

#### CASC5

Forward 5’-TCGCTGAAGTGGAAACAGAAAC-3’

Reverse 5’-TATCTGAGCAAGGGTCTCTGCG-3’

#### WDR62

Forward 5’-GCTGACAAATGGCAAGCTG-3’

Reverse 5’-GATGGTCTTGAGGGGTTCCT-3’

### Hematoxylin & Eosin

After three weeks of age, pups were sacrificed, and brain tissues removed postmortem were embedded in optimal cutting temperature (OCT) compound. Cryostat sectioned slices of 5-micron thickness were stained with hematoxylin and eosin (H&E) as described previously (16). Images were acquired using an inverted fluorescence microscope with a 560 Axiovision camera and 20X and 40X objectives (Zeiss, Germany).

### Murine mixed glial cell culture

For primary murine glial culture, postnatal day 4-6 (P4-P6) Atg6^+/−^ and Atg6^+/+^ littermates were separated according to phenotypic coat color and sacrificed according to IACUC guidelines as described previously (17, 18). Cells seeded in 6 well plates were infected with ZIKV at MOI of 0.1 or treated with ZIKV envelope (E) and the non-structural protein (NS)-1 proteins. Viral proteins were purchased from ImmunoDx, Woburn, MA, USA. The protein concentration used (50nM) was based on a dose response curve and concentrations reported in cerebral spinal fluid (CSF) of patients with flavivirus infection (19). Since viral proteins were resuspended in PBS after purification with aqueous solvent, PBS was used as a negative control.

### ELISA

Cell culture supernatants (pre-cleared by brief centrifugation) were used to measure the levels of interleukin (IL)-6, monocyte chemotactic protein-1 (MCP-1), regulated on activation, normal T cell expressed and secreted (RANTES) and tumor necrosis factor alpha (TNF-α) by ELISA (R&D Systems, Minneapolis, MN, USA) according to the manufacturer’s instructions. The optical density (O.D.) was read at A450 on a Synergy HTX plate reader (BioTek, Winooski, VT, USA).

### Plaque Assay

Vero cells were infected with a 10-fold dilution of ZIKV stock or the supernatants from infected/treated cells. After 1-hour adsorption cells were washed with PBS. Cells were overlaid with culture media (EMEM supplemented with 2% FBS) containing equal volume of 3.2% carboxymethylcellulose and incubated for 5 days at 37°C. Cells were fixed and stained with 1% crystal violet solution prepared in 20% formaldehyde, 30% ethanol and 50% PBS for 1 hour. Stained cells were washed with water to remove excess crystal violet, left to dry overnight, and lysis plaques were quantified by stereomicroscope (Zeiss). The viral titer was expressed as plaque forming units (PFU) per ml of the stock (13)

### Immunohistochemistry

ZIKV infectivity was measured by fluorescent immunolabeling as described by Ojha et al. (13). Briefly, cells and brain tissue sections were fixed in 4% paraformaldehyde, permeabilized with 0.1% Triton X-100, and blocked in 10% milk/0.1% goat serum. Sections were immunolabeled with the neuronal marker, mouse anti-MAP2 (microtubules associated protein 2) antibody (Cat. MAB378, Millipore, Boston, MA, USA), anti-ZIKV-E antibody (Cat. GTX133314) and anti-ZIKV-NS1 antibody (Cat. GTX133307, Genetex, CA, USA). Immunoreactivity was visualized with secondary antibodies from Molecular Probes (Carlsbad, CA, USA). 4′,6-diamidino-2-phenylindole (DAPI) was used to label cell nuclei. Images were analyzed using an inverted fluorescence microscope with a 560 Axiovision camera (Zeiss).

### Western Blotting

Protein was extracted from postmortem brain tissues of both Atg6^+/+^ and Atg6^+/−^ animals using RIPA buffer (Thermo Scientific, Waltham, MA, USA) supplemented with a mixture of protease and phosphatase inhibitors followed by SDS-PAGE protein separation. Immunoblots were labeled with primary antibodies against Beclin1 – 1:500 (Novus Biologicals, NB500–249), ATG5 – 1:200 (Novus Biologicals, NB110-53818), LC3-B – 1:1000 (Novus Biologicals, NB600–1384), P62/SQSTM1– 1:500 (Novus Biologicals, NBP1–48320) and β actin (Cat. sc-47778, 1:200) (Santa Cruz Biotechnology, Santa Cruz, CA, USA) was used as internal control. Immunoblots were subsequently incubated with secondary antibodies conjugated to horseradish peroxidase (Millipore, Billerica, MA, USA), exposed to SuperSignal West Femto Substrate (Thermo Scientific) and visualized using a ChemiDoc imaging system (Bio-Rad,). Densitometric analysis was quantitatively measured using image J (NIH.gov).

### Statistical analysis

Results are reported as mean ± SEM of 3-5 independent experiments. Data were analyzed using analysis of variance (ANOVA) followed by the post hoc test for multiple comparisons (GraphPad Software, Inc., La Jolla, CA, USA). An alpha level (p-value) of < 0.05 was considered significant.

## RESULTS

### Pregnant Atg6^+/+^ and Atg6^+/−^ dams transiently immunosuppressed are susceptible to ZIKV infection

We explored the role of Beclin1 in ZIKV infection and disease using timed-pregnant Beclin1 deficient (Atg6^+/−^) and wild-type (Atg6^+/+^) mice model. For the *in vivo* studies, genotype of each animal strain was confirmed by PCR (15) followed by detection of protein expression levels by western blotting (Figure 1). Representative immunoblots confirmed a decrease in Beclin1 and LC3-II expression levels and increased in p62/SQSTM1 levels in tissues extracted from Atg6^+/−^ mice when compared to Atg6^+/+^ mice (Fig. 1A and B). For ZIKV infection, pregnant dams received anti-interferon receptor 1 (anti-IFNAR1) monoclonal antibody at 2mg/animal via intraperitoneal (ip) route at gestational day 8 followed by subcutaneous (sc) infection with individual strain of ZIKV at 10^3^ plaque forming unit (PFU) in 50µL of PBS or mock (PBS) injection at gestational day of E9. To confirm infection, serum was removed at gestation day E13 (4 days post-infection) and viral RNA was measured by RT-PCR. Viral RNA levels in the range of 10^3^ PFU/mL were detected in serum of Atg6^+/+^ and Atg6^+/−^ dams infected with ZIKV-R103451 and ZIKV-MR766, while infection with ZIKV-PRVABC59 showed significantly lower viral RNA levels in serum of Atg6^+/−^ dams when compared to Atg6^+/+^ dams (Fig. 1D). The weight of each pregnant animal was measured before the detection of a vaginal plug and throughout the gestation period. Increases in body weight served as a measurable indicator of pregnancy among dams (Fig. 1E). As expected, Beclin1 deficient (Atg6^+/−^) animals showed less gain in body weight compared to wild-type (Atg6^+/+^) dams. This was likely because Atg6^+/−^ delivered fewer pups when compared to Atg6^+/+^ dams, since litter numbers delivered by Atg6^+/−^ dams (crossed with an Atg6^+/−^ sire) are controlled by their genetic background. Within a litter, approximately 50% of the pups delivered are heterozygous for the beclin1 gene (Atg6^+/−^). These animals have an agouti coat color which is believed to be a result of the effect of the Becn1 mutation on melanogenesis. 25% of the pups delivered are homozygous for the beclin1 gene (Atg6^+/+^) with a black coat color, while homozygous deletion of the targeted allele results in embryonic lethality (~25%). Litter numbers ranged between 5 to 7 pups for Atg6^+/−^ and between 6 to 9 pups for Atg6^+/+^ per litter. Linear regression models (based on weight change from day (0) demonstrated that maternal weight gain at day 11 was a significant predictor of litter size (Fig. 1E). At E17 (8 days post-infection), maternal placenta and other organs removed postmortem from Atg6^+/+^ and Atg6^+/−^ dams, showed a high level of viral RNA in the placenta, followed by the spleen, liver, heart and the lowest titer was detected in the brain, irrespective of mice strain. The low level of viral RNA detected in the brain is indicative that ZIKV can cross the blood brain barrier (Fig. 1F). The survival rates in pregnant dams’ post-infection with ZIKV was also monitored for the duration of gestation and showed no significant differences between mock (PBS) and ZIKV-R103451 infected Atg6^+/+^ dams when compared to similar treated Atg6^+/−^ dams (Fig. 1G). On the contrary, increased fatality was observed in animals infected with ZIKV-MR766 and to lesser extend in Atg6^+/−^ dams infected with ZIKV-PRVABC59 (Fig. 1G). Despite the high mortality rate detected in dams infected with ZIKV-MR766 when compared to dams infected with ZIKV-R103451, similar levels of viral RNA were detected in serum and organs recovered at E17, suggesting that different viral isolates exhibit different pathogenicity. Because of the high mortality among dams infected with ZIKV-MR766, we sought to confirm whether the observed mortality was caused by ZIKV or because of animal strain. Infection using AG129 mice lacking receptors for both Type I and Type 2 IFN, infected with increasing doses (10^1^, 10^2^, 10^3^ and 10^4^ PFU/ml) of ZIKV-MR766 showed an infection-dose dependent decrease in survival rate of the mice, confirming that mortality was probably due to viral infection and not necessarily due to mouse genotype. (Fig. 1H). Overall, the data shows significant infection throughout gestation in both Beclin1 (Atg6^+/−^) deficient and wild-type (Atg6^+/+^) pregnant dams infected with three different strains of virus. The data also shows that among the three isolates, ZIKV-MR766 is more lethal irrespective of mouse genotype.

**Figure 1.**
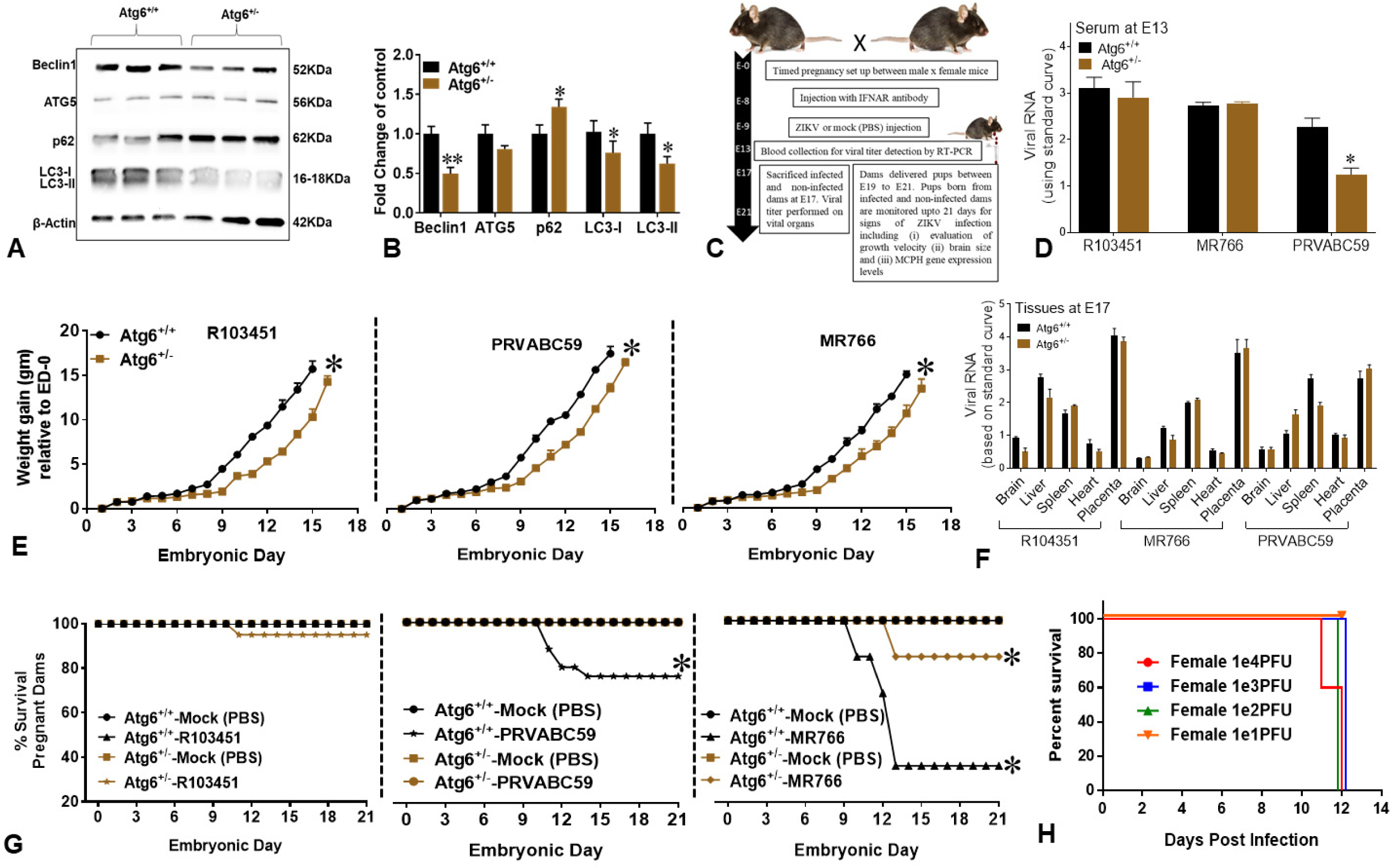
ZIKV infection in Atg6^+/+^ and Atg6^+/−^ pregnant dams. (A) Representative western blots probed with antibodies against several autophagy proteins. Adult Atg6^+/+^ and Atg6^+/−^ brains were removed postmortem and minced according to Materials and Methods. (B) Densitometric analysis using image J indicate the levels of Beclin1, ATG5, P62, LC3-I and LC3-II in brains of adult Atg6^+/+^ (black bar) and Atg6^+/−^ (brown bar) mice. \**p*<0.05 vs. Atg6^+/+^. (C) Schematic diagram illustrating ZIKV-infection in timed-pregnant dams. (D) Viral RNA detected in serum collected from ZIKV-infected dams on E13. \**p*<0.05 vs. Atg6^+/+^. (E) Weight gain, expressed in grams, was measured using an analytical balance at gestation day 0 and throughout pregnancy at 3 days interval. \**p*<0.05 vs. Atg6^+/+^. (F) Viral RNA detected in organs removed postmortem from ZIKV-infected dams on E17. (G) Percent survival rate in pregnant dams infected with ZIKV or mock (PBS) was calculated by dividing the total number of live animals by the number of live + dead animals X 100. No significance difference (p = 0.179) in R103451-infected dams. \**p*<0.05 vs. mock exposed. (H) Percent survival in adult AG129 mice infected with ZIKV-MR766 (10^1^ to 10^4^ PFU/mL). Error bars show mean ± SEM for N= 5 - 8 animals per treatment. The data were analyzed using GraphPad Prism and two-way ANOVA followed by Tukey’s test. * indicates p<0.05 and ** indicates p<0.01. Viral RNA is expressed on a log10 scale after comparison with a standard curve produced using serial 10-fold dilutions of ZIKV RNA from known quantities of infectious virus.

### Growth impairment in pups exposed to ZIKV-R103451 in utero

Subcutaneous infections in pregnant dams with ZIKV on E9 and embryo harvested after eight days on E17 showed the vertical transmission of ZIKV from the placenta to the fetus (Fig. 2A). It is important to note that at E17, a period of neurogenesis, no noticeable growth abnormality was observed in fetuses, irrespective of animal strains or viral phylogeny (data not shown). At E20-E21, pups born to mock (PBS) and ZIKV-infected Atg6^+/+^ and Atg6^+/−^ dams were monitored for up to 21-days for mortality and for morphological abnormalities. A slight decrease in the survival rate of pups born to ZIKV-R103451-infected Atg6^+/+^ and Atg6^+/−^ dams was noted when compared to mock-infected animals (Fig. 2B), while the survival rate in pups born to ZIKV-PRVABC59 and ZIKV-MR766-infected dams were considerably low (Fig. 2B). In fact, greater than 80% of pups born to ZIKV-MR766-infected dams died after 2 days postnatal (Fig. 2B; middle panel). In Fig 2C (top panel) is illustrated a representative image of a litter born to ZIKV-R103451-infected Atg6^+/−^ dam that consisted of both Atg6^+/+^ (black) and Atg6^+/−^ (agouti) pups. Within a litter, the smaller pup is indicated by a circle at day 7 and with an arrow at day 10. Fourteen days post-birth, growth abnormalities became exceedingly visible by differences in body size and was detected in 1 of every 4 (25-30%) pups. Genotyping confirmed that the small pups were heterozygous for the *beclin1* gene (data not shown). The average body weight was approximately 6.93 gm (Fig. 2C; top chart) and the average body length was around 5.38 cm (Fig. 2C, bottom chart). After 21 days, both small and typical sized pups were sacrificed, and brains were removed for further analysis. Representative images of 3-week-old pups born from ZIKV-R103451-infected (top) and mock (bottom) infected dams are shown in Fig. 2D. Respective skull and brain images are shown on the right-hand side. Brain recovered from the small pups born to ZIKV-R103451-infected dams are labeled 3 and 4, while brain recovered from the typical sized pups born to ZIKV-R103451-infected dams are labeled 1 and 2. Brain recovered from the typical sized pups born to mock-infected dams are labeled 5-8. The weight (in milligram) of each brain determined by a balance is represented in a bar graph (Fig. 2E: top) and in a chart (Fig. 2E: bottom). As expected, the smaller brains weigh less than the well-defined brains. Except for one pup born to a ZIKV-PRVABC59-infected dam, no significant growth abnormalities were measured in pups born to dams infected with ZIKV-PRVABC59 or ZIKV-MR766, irrespective of murine strains (data not shown). Overall, the data shows a growth dysfunction in pups born to ZIKV-R103451 that was reflected in body weight, body length and brain weight parameters. The impairment in body growth along with the abnormal brain morphology was higher among pups born to Atg6^+/−^ mice infected with ZIKV-R103451, suggesting a potential function of Beclin1 in growth development.

**Figure 2.**
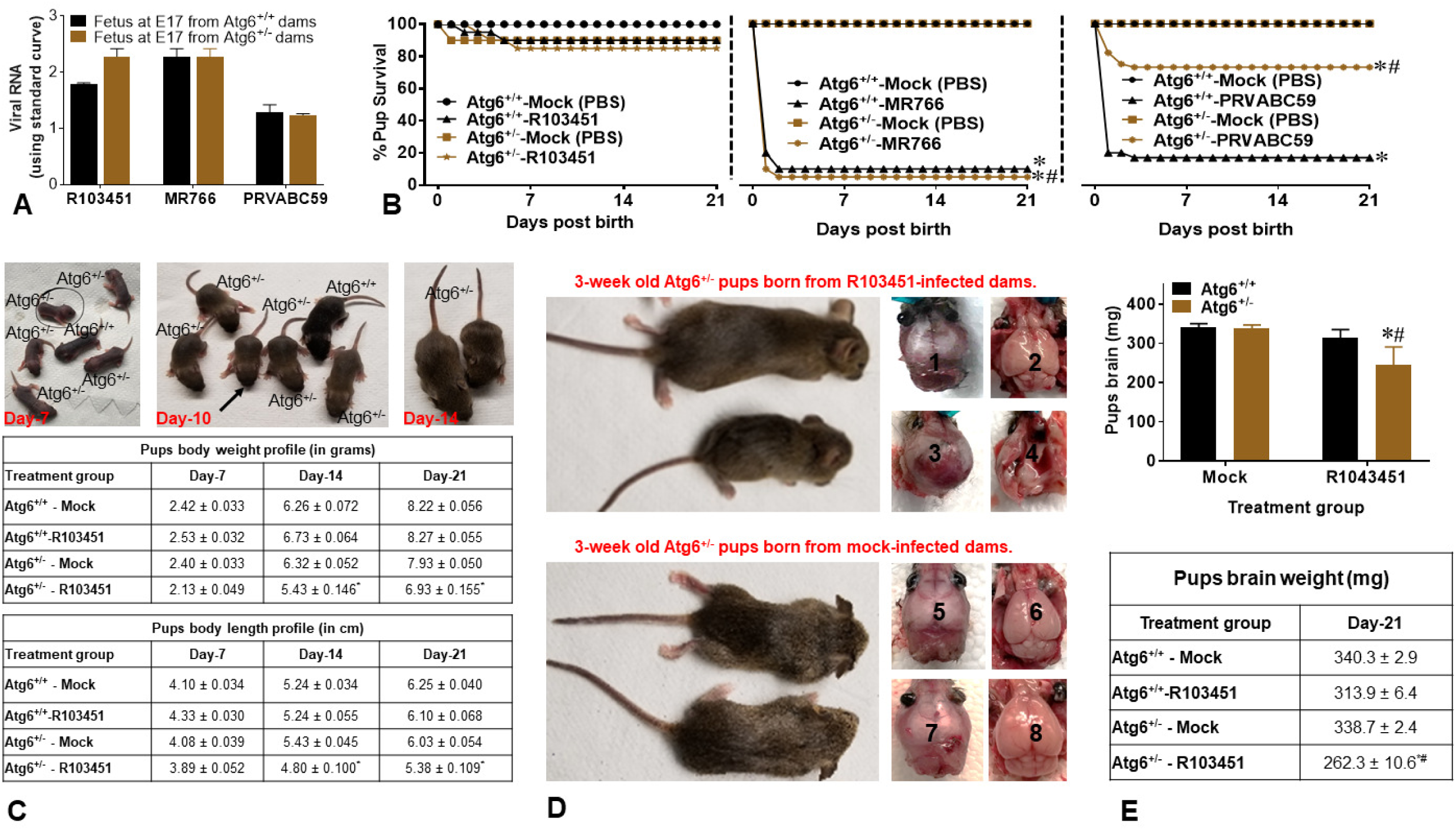
Growth impairment in pups born to ZIKV-R103451-infected dams. (A) Viral RNA was measured in postmortem fetuses collected on E17 from Atg6^+/+^ and Atg6^+/−^ dams infected with ZIKV. (B) Percent survival rate in pups born to ZIKV-infected dams was calculated using the total number animals subtracted by the dead pups X 100. (C) Representative image of a litter born to ZIKV-R103451 Atg6^+/−^ dam containing both Atg6^+/+^ and Atg6^+/−^ pups. Smaller pup is shown in a circle at day 7, with an arrow at day 10, and at day 14, the smaller sized pup becomes more noticeable when compared to the regular sized sibling (Top panel). Body weight (middle chart) and body length (bottom chart) profile of Atg6^+/+^ and Atg6^+/−^ pups born from mock and ZIKV infected dams. Body weight was measured using a balance and expressed in grams while body length was measured using a caliper and expressed in centimeter. (D) Representative images of 3-week-old pups born from ZIKV-R103451-infected (top) and mock (bottom) infected dams are shown in Fig. 2D. Respective skull and brain images are shown on the right-hand side. Brain recovered from the small pups born to ZIKV-R103451-infected dams are labeled 3 and 4, while brain recovered from the typical sized pups born to ZIKV-R103451-infected dams are labeled 1 and 2. Brain recovered from the typical sized pups born to mock-infected dams are labeled 5-8. (E) Brains weight in milligrams are represented in bar graph (top) and in a chart (bottom). Error bars show mean ± SEM for N = 64 (Atg6^+/+^) and N = 48 (Atg6^+/−^) pups used. The data were analyzed using Two-way ANOVA followed by Tukey’s test for multiple comparison. \**p*<0.05 vs. mock-infected Atg6^+/+^ strain, whereas, #*p*<0.05 vs. mock infected Atg6^+/−^ strain. Viral RNA is expressed on a log10 scale after comparison with a standard curve produced using serial 10-fold dilutions of ZIKV RNA from known quantities of infectious virus.

### Dysregulation of autophagy exacerbates the pathology in pups exposed to ZIKV-R103451 in utero

Potential causal factor(s) responsible for the growth impairment detected in pups born to ZIKV-R103451-infected Atg6^+/−^ dams was further explored. After three weeks, pups born to ZIKV-infected or mock-exposed dams were sacrificed and brains removed postmortem were snap-frozen in liquid nitrogen for further use. Half of the brain hemisphere was used for ELISA and RT-PCR while the other half of the brain was used for imaging analysis. Sections of the frontal cortex from brain recovered postmortem in Atg6^+/−^ pup were used for immunofluorescent imaging (Fig. 3A); similar staining pattern were detected in sections closer to the center of the brain (data not shown). Although viral RNA was not detected by RT-PCR (data not shown), immunofluorescent double labeling with antibodies against the neuronal marker, MAP2 (labeled in red) and ZIKV proteins (labeled in green) showed expression of NS1 (Fig. 3A; left bottom panel) and the structural E protein (Fig. 3A; right bottom) in brain tissues recovered from Atg6^+/−^ pups born to ZIKV-infected dams. Brain tissues recovered from Atg6^+/−^ pups born to mock-exposed dams showed no fluorescent labeling with NS1 or E antibodies (Fig. 3A; top panels). Microscopic appearance of brain stained with H&E showed no visible sign of aberrant morphology of neurons in mock-exposed brain tissues removed postmortem (Fig. 3B; left panel). In contrast, brain tissues recovered from Atg6^+/−^ pups born to ZIKV-infected dams showed signs of necrotic neurons with shrunken neuronal cell bodies (Fig. 3B; right panel). The other half of the brain hemisphere was minced and used to measure the expression of autophagy-related genes and growth factors, crucial for neurodevelopment and homeostasis, by RT-PCR. The insulin-like growth factor-1 (IGF-1), a polypeptide hormone with critical roles in regulating brain plasticity mechanisms, was reduced by 8-fold in Atg6^+/−^ pups born to ZIKV-infected dams when compared to a 4-fold decrease in Atg6^+/+^ pups born to ZIKV-infected dams (Fig. 3C), suggesting a potential link between IGF-1 and ZIKV-associated growth impairments. The transmembrane protein 74 (TMEM74), a novel autophagy-related protein, was upregulated by approximately 6-fold in Atg6^+/−^ pups born to ZIKV-infected dams. TMEM74-related autophagy is independent of BECN1/PI3KC3 complex, which may explain the reason this gene was more expressed in animals lacking the Beclin1 gene and also in the context of our animal model may not be linked to ZIKV exposure (20). Additional genes involved in the autophagy machinery are also included in the graph, although no significant differences were detected between Atg6^+/+^ and Atg6^+/−^ pups (Fig. 3C). Expression levels of several microcephaly-related genes previously linked to stillbirth, brain development, and microcephaly in fetuses, were also measured by RT-PCR (8, 21–23). Gene expression levels of MCPH1 and ASPM in brain tissues of Atg6^+/−^ pups born to mock (PBS)-exposed dams were significantly reduced when compared to Atg6^+/+^ pups born to mock-exposed dams (Fig. 3D; top graphs); implying an important role of Beclin1 in growth development. Likewise, expression levels of MCPH1, ASPM, CASC5 and WDR62 in brain tissues of Atg6^+/−^ pups born to ZIKV-R103451 infected dams were significantly lower (approximately 2.5 - 3-fold) when compared to Atg6^+/+^ pups born to ZIKV-R103451-infected dams (Fig. 3D). Surprisingly, RNA expression levels of the microcephalic genes MCPH1, ASPM, CASC5, and WDR62 were also decreased in brains removed from Atg6^+/+^ and Atg6^+/−^ pups born to ZIKV-MR766 and ZIKV-PRVABC59-infected dams (data not shown), despite no growth abnormalities among these pups, suggesting that overt growth impairment detected in ZIKV-R103451 exposed pups may not be exclusively linked to changes in microcephalic genes. Overall, the data shows a decrease in the expression of growth factors with visible signs of necrotic neurons in brain recovered from Atg6^+/−^ pups; these factors may or may not be associated with the observed morphological abnormalities.

**Figure 3.**
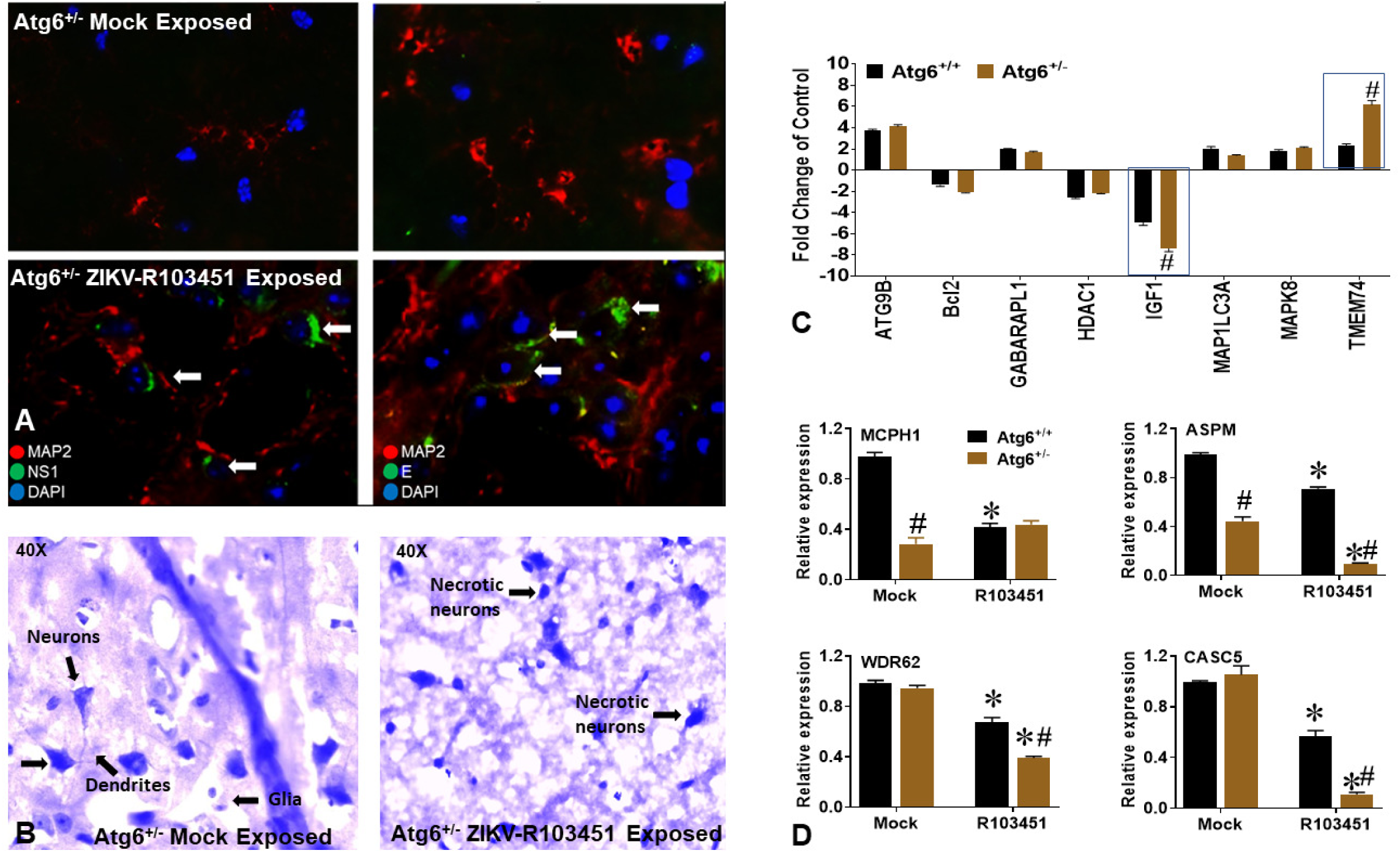
Reduced expression of microcephalic genes in brain of pups exposed *in-utero* to ZIKV. (A) Three weeks post-birth, pups born to ZIKV-infected and mock-exposed dams were sacrificed and brains removed postmortem were embedded in OCT and used for imaging analysis. Representative images of brain tissues from pups exposed to mock (top panel) and ZIKV (bottom panel) *in utero*, labeled with MAP2 expressing neurons (indicated with red fluorescent color), NS1 (left) and E (right) (arrows: indicated with green fluorescent color) and the blue color indicates DAPI-labeled nuclei. (B) Hematoxylin & Eosin (H&E) staining of brains removed postmortem from pups exposed to mock (left panel) and ZIKV (right panel) *in utero*. Images were acquired using an inverted fluorescence microscope with a 560 Axiovision camera using 40X magnification (Zeiss). Neurons, dendrites, glia and necrosis are indicated by arrows. (C) The other half of the brain tissues were minced and used to measure autophagy related genes and growth factors in Atg6^+/+^ (black bar) and Atg6^+/−^ (brown bar) pups exposed to ZIKV-R103451 in *utero*, by RT-PCR. Results are expressed as fold change from control (increase or decrease). (D) RNA expression of the microcephaly associated genes (MCPH1, ASPM, WDR62, and CASC5) were measured by RT-PCR in Atg6^+/+^ (black bar) and Atg6^+/−^ (brown bar) pups exposed to mock and ZIKV-R103451 in *utero*. Expression levels are relative to Atg6^+/+^ pups born from wild type mice and normalized to GAPDH. Error bars show mean ± SEM for N = 64 (Atg6^+/+^), N = 48 (Atg6^+/−^) pups. The data were analyzed by Two-way ANOVA followed by Tukey’s multiple comparison test. \**p*<0.05 vs. respective mock infected strain, ^#^*p*<0.05 vs. Atg6^+/+^

### Beclin1 deficiency exacerbates secretion of inflammatory molecules in ZIKV-infected glia in vitro

Since glial cells are the most abundant cell types in the brain and the principal cell types involved in the release of neuroinflammatory molecules, they are frequently considered the culprit in many viral pathologies (24–26). To this end, mixed glia (astrocytes and microglia) isolated from whole brain of either Atg6^+/+^ or Atg6^+/−^ pups, as described previously (15), were infected with ZIKV at an MOI of 0.1. Mixed glial cultures were permissive to infection with ZIKV (R103451, PRVABC59, MR766), albeit more level of infection was detected in Atg6^+/−^ glia infected with ZIKV-R103451. Fig. 4A, shows a representative image of glia derived from Atg6^+/+^ and Atg6^+/−^ pups infected with ZIKV after 24-hours, followed by immunofluorescent labeling with the antibody against GFAP (red), ZIKV NS1 (green) and DAPI nucleus (blue). Viral infection and PFU were analyzed by plaque assays (Fig. 4B: top panel) using supernatants collected at various time-points post-infection (Fig. 4B; bottom panel). Next, the secretion of inflammatory molecules was measured by ELISA using supernatant from non-infected (media) and ZIKV-infected glia. Infection with ZIKV-MR766 and ZIKV-R103451 and to a lesser extend ZIKV-PRVABC59 caused a significant increase in RANTES, MCP-1 and IL-6 at 24-hours that was still detected after 48-and 72-hours post-infection (Fig. 4C). At twenty-four-hours post-infection with ZIKV-R103451, secretion of RANTES was increased by a 2.5-fold, MCP-1 was increased by a 1.4-fold, and IL-6 was increased by 1.6-fold in supernatant derived from Atg6^+/−^ infected glial cells when compared to glia derived from Atg6^+/+^ pups (Fig. 4C). More importantly, the secretion of MCP-1 in supernatant from Atg6^+/−^ glia infected with ZIKV-R103451 remained high throughout the duration of the experiment (Fig. 4C). Since we were able to detect NS1 and E proteins, in the absence of viral RNA, we posit that expression of these proteins might be the contributing components toward neuroinflammation. Cultured glia cells were incubated with 50nM of recombinantly expressed NS1 and E proteins. This concentration was based on a dose response curve (data not shown) and concentration of proteins reported in the cerebral spinal fluid of patients with flavivirus infection (19) and in sera of DENV-infected patients (27, 28). As expected, direct exposure of murine glia to NS1 or the E protein caused a significant secretion in inflammatory molecules, with the most pronounced effects observed with TNF-α expression in supernatant from Atg6^+/−^ glia (Fig. 4D). Overall, the data shows infection of ZIKV in murine-derived glia along with the secretion of inflammatory molecules. Furthermore, high level of viral RNA measured in ZIKV-infected Atg6^+/−^ glia after 24 and 48-hours post-infection correlates with the increased secretion of inflammatory cytokines. Findings also point to a potential link between Beclin1 and the regulation of TNF-α, in response to pathogenic insults, which may also account for the phenotype detected in pups born to ZIKV infected Atg6^+/−^ dams.

**Figure 4.**
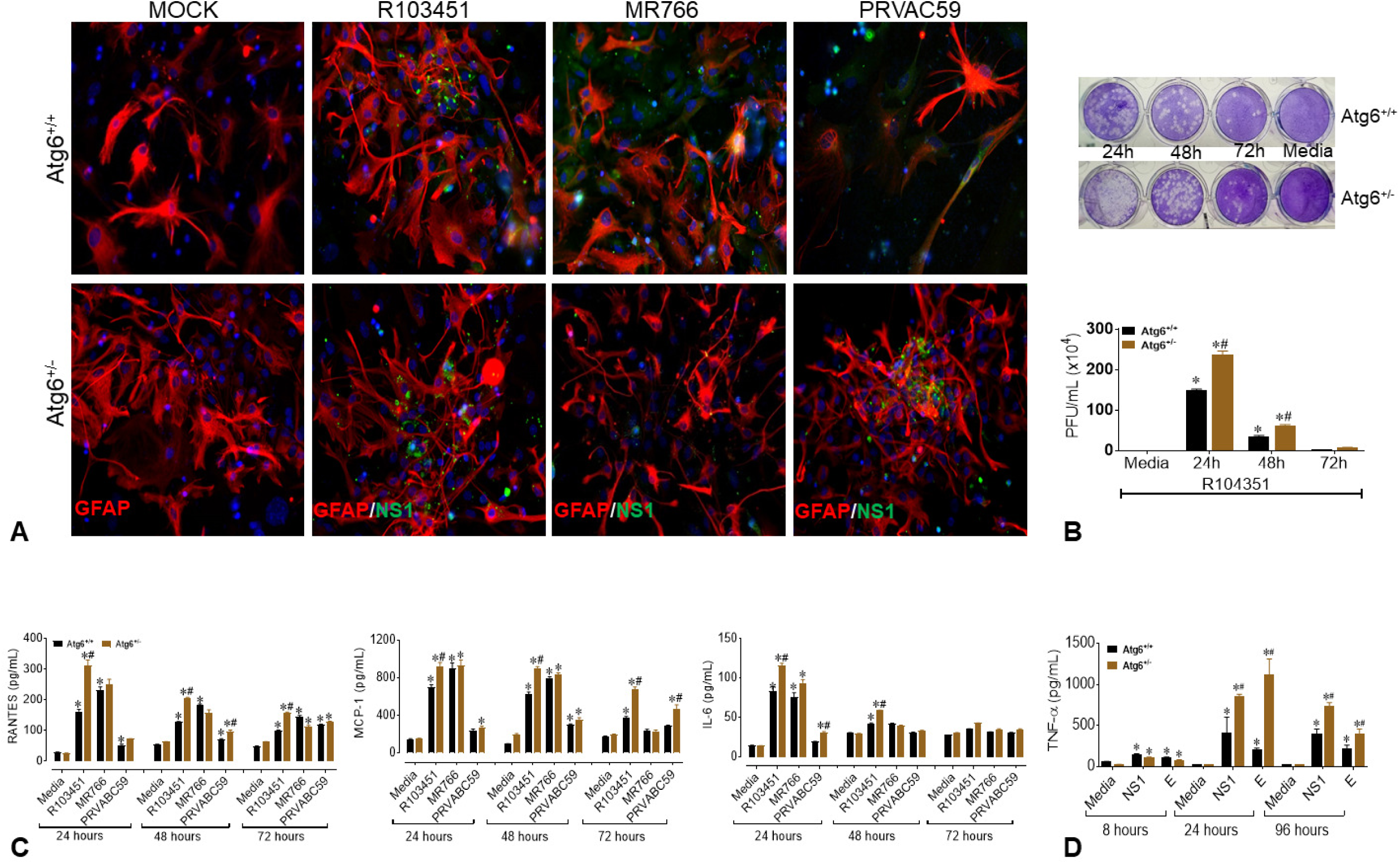
ZIKV infects mixed mouse glia and induces inflammatory molecules. (A) Representative immunofluorescent images of mouse mixed glia derived from Atg6^+/+^ and Atg6^+/−^ pups infected ZIKV and labeled with the antibody against GFAP (red), ZIKV NS1 (green) and DAPI nucleus (blue). Images were acquired using an inverted fluorescence microscope with a 560 Axiovision camera and a 40X magnification (Zeiss). (B) Viral infection and PFU were analyzed using plaque assays (top) and data are illustrated in graph (bottom). Atg6^+/+^ glia (black bar) and Atg6^+/−^ glia (brown bar). (C) Secretion of RANTES, MCP-1 and IL-6 were detected in glial supernatants infected with ZIKV at 24, 48- and 72-hours post-infection by ELISA. (D) Secretion of TNF-α was detected in glial supernatants exposed to 50nM of viral proteins after 8, 24- and 96-hours by ELISA. Atg6^+/+^ glia (black bar) and Atg6^+/−^ glia (brown bar). Error bars show mean ± SEM for 3 independent experiments. The data were analyzed by Two-way ANOVA followed by Tukey’s multiple comparison test. \**p*<0.05 vs. respective media control, ^#^*p*<0.05 vs. Atg6^+/+^.

## DISCUSSION

In this study, we reported for the first time that three different phylogenetic strains of ZIKV infects timed-pregnant Beclin1-deficient (Atg6^+/−^) and wild-type (Atg6^+/+^) mouse model (Fig. 1). Impact of ZIKV infection on dams were detected at E13 in serum and at E17 in placenta and in other organs removed postmortem with limited viral RNA detected in the brain, despite the use of anti-IFNAR1 mAb (Fig. 1). Low RNA detection in the brain is not unusual, since a report by Cao et. al 2017, also reported low levels of viral titers (in the range of 10-100 FFU equivalent/g) in fetal brain (29). Except for one pup born to a ZIKV-PRVABC59-infected dam, no significant growth abnormalities were measured in pups born to dams infected with ZIKV-PRVABC59 irrespective of murine strains. This finding was unexpected, since the Honduran and the Puerto Rican strains of ZIKV arose from the same 2015 outbreak. Placenta recovered from postmortem dams infected with the Honduran strain of ZIKV showed higher viral RNA levels when compared to the placenta recovered from the ZIKV-PRVABC59-infected dams (Fig. 1). The low level of placental infection detected in the PRVABC59-infected dams could have contributed to the lack of phenotypic abnormalities detected in the pups. On the contrary, 30% of pups heterozygous for the Atg6 gene born to ZIKV-R103451-infected dams showed growth impairment (Fig. 2). No evidence of viral RNA was detected in 3-week-old pups, despite evidence of growth impairment (Fig. 2). A lack in viral RNA detection may indicate the absence of virions or low RNA detection limit by the RT-PCR but also, it should be reminded that pups did not receive IFNAR mAb postnatal and this could account for the lack of viral detection, as the normal immune system of the pups may have effectively suppressed viral RNA below the limit of quantification. It is not unexpected that only viral proteins, but not viral RNA was detected. In fact, in a panel of patient sera infected with DENV, the NS1 protein was detected even in the absence of viral RNA or in the presence of immunoglobulin M antibodies. NS1 circulation levels varied among individuals during the course of the disease, ranging from several ng/mL to several ug/mL (27). Likewise, presence of viral protein in the absence of viral RNA was reported in serum recovered from HIV-positive subjects treated with antiretroviral drugs, implying that viral RNA can be suppressed below detection level, while maintaining detectable protein expression in leaky reservoirs (30).

Necrotic neurons were detected in sections of the frontal cortex area in postmortem brain recovered from 3-week-old pups. Although mechanism(s) mediating growth impairments with ZIKV infection are still unclear, viral infection itself can damage neural progenitor cells or alternatively, ZIKV mediated reduction in the expression of microcephaly related genes which are directly involved in neuronal cell division and proliferation may also contribute to the impairment in brain development (23, 31). Others have shown that mutations in the human *WDR62* resulted in microcephaly and a wide spectrum of cortical abnormalities (32–34), while a loss in the WDR62 protein function in mice causes mitotic delay, death of neuron progenitor cells, reduced brain size and dwarfism (34). *CASC5* was shown to be involved in cell cycle and kinetochore formation during metaphase with mutation in this gene was also implicated in causing microcephaly (35). Using a mouse models of *Mcph1* mutations it was shown that microcephaly can develop due to premature differentiation of neurons (50). Likewise, gliosis and neuronal necrosis were previously associated with ZIKV infected microcephalic brain (51, 52). Thus, one possible explanation for the observed morphological changes in neurons could be related to the decreased expression in microcephalic genes, while attenuated Beclin1 expression further exacerbated the pathology. However, a decrease in the expression of microcephalic genes was also detected in brains of Atg6^+/+^ pups born to ZIKV-R103451 infected dams as well as in brains of Atg6^+/+^ and Atg6^+/−^ pups born to ZIKV-MR766 and ZIKV-PRVABC59 infected dams, despite no detection of necrotic neurons (data not shown). A significant downregulation in the expression of the microcephaly related genes, MCPH1 and ASPM, in the brains of Atg6^+/−^ but not in Atg6^+/+^ pups born to mock-exposed dams while expression levels of MCPH1, ASPM, WDR62, and CASC5 were reduced in the brains of Atg6^+/−^ and Atg6^+/+^ pups born to ZIKV-exposed dams, irrespective of viral strain (Fig. 3). Reduction in Beclin1 or impaired autophagy enhanced ZIKV-R103451 (but not other strains)-mediated pathology in *in-utero* exposed pups; suggesting a ZIKV strain specific effect of autophagy pathway in associated pathologies. Beclin1 and the ultraviolet irradiation resistance-associated gene (UVRAG) are involved in both autophagy and centrosome stability and linked to ZIKV mediated microcephaly (36, 37), while the recently identified MCPH18, a phosphatidylinositol 3-phosphate-binding protein, functions as a scaffold protein for autophagic removal of aggregated protein; suggesting a potential link of autophagy in the development of primary microcephaly (38). Autophagy is a common pathway involved in regulating the replication of ZIKV as well as other viral-infections in cells of the central nervous system (13, 17, 39–43). In a related studies published by others, an autophagy-deficient animal model lacking the Atg16L gene showed restricted ZIKV infection in placenta, with reduced ZIKV-mediated placental damage and reduced adverse fetal outcomes (44). Reduction of ATG16L1 expression levels in pregnant dams or placental trophoblastic cells showed limited ZIKV burden which contradicts our current studies, as we did not detect a decrease in ZIKV infection in dams using an autophagy-deficient animal model heterozygous for the Atg6 gene. Although speculative, the discrepancy between ATG16L1 and ATG6 knock down (used in our studies) may relate to the differential role of the specific protein in the autophagy pathway and how specific steps in autophagy influence the life cycle and pathology of ZIKV.

As for our findings, further studies including gene silencing and protein overexpression are needed to better understand and decipher the cause and effect of the microcephalic genes in our animal model. Alternatively, the low expression of IGF-1 detected in postmortem brains of pups heterozygous for the Atg6 gene born to ZIKV-R103451-infected dams may have triggered neuronal injury and subsequently downregulated the microcephalic genes. The IGF system plays a central role in hormonal growth regulation and is responsible for normal fetal and postnatal growth. For more than 30-years, IGF has been available as a replacement therapy in growth hormone-deficient patients and for the stimulation of growth in patients with short stature of various causes (45). In a case study, a disruption of the IGF system in patient was associated with microcephaly, growth retardation, and intellectual disability (46). Using a mice model with IGF-1 gene knockout, animal presented with microcephaly and demyelination in the whole brain (47), whereas overexpression of IGF-1 was shown to cause macrocephaly. The concentrations of IGF-1 in the cerebral spinal fluid have been correlated with brain growth in autistic children (48, 49) while low values of IGF-1 have been reported in a number of serious neurologic diseases of children (50). Since levels of IGF-1 was significantly reduced in Atg6^+/−^ brain recovered in pups at 3 weeks of age (Fig. 3C), this may be another underlying factor associated with the phenotype detected in our *in vivo* infectious model while autophagy is required for proper functionality (51, 52).

As noted above, a lack of viral RNA was detected in postmortem brains recovered in 3-week-old pups, which led to subsequent *in vitro* studies, to determine the significance of the secreted proteins in the pathology of ZIKV. Presence of viral proteins in the central nervous system can cause neuroinflammation, glial dysfunction, excitotoxicity, and neuronal death (17, 53). Glia have been found to play key roles in neuroinflammation, and although this is a normal and necessary process, emerging evidence in animal models suggests that sustained inflammatory responses by glia can contribute to disease progression (54, 55) and possibly considered as a general underlying factor associated with the phenotype detected in our *in vivo* infectious model. Although the *in vitro* data does not necessarily support the causal factors detected in the *in vivo* studies, the *in vitro* data does confirm that our mouse model can be infected with three isolates of ZIKV and that attenuated Beclin1 is associated with an increase in viral replication (Fig. 4B) which correlated with an increase in viral-induced chemokine (Fig. 4C) and viral protein-induced cytokine secretion (Fig. 4D).

In summary, we showed (i) infectivity of three different ZIKV isotypes using a conventional mouse model and (ii) we showed growth impairment in Beclin1-deficient pups exposed to ZKIV-R103451 *in utero* without detection of viral RNA in pups; suggesting that while ZIKV itself can cause disease there are other factors and probably an indirect role of Beclin1 and the autophagy pathway associated in ZIKV infection and pathology.

## ACKNOWLEDGEMENTS

Dr. Nazira El-Hage received funding from the Florida Department of Health (http://www.floridahealth.gov/) award number 7ZK09 and the National Institutes of Health (https://www.nih.gov/) grant number DA036154. The funders had no role in study design, data collection and analysis, decision to publish, or preparation of the manuscript. We acknowledge Mr. Hary Estrada-Bueno (Senior Laboratory Technician) who assisted with the timed pregnancy set-up. We also acknowledge the university graduate school of FIU for the presidential fellowship and the dissertation year fellowship provided to Mr. Chet Raj Ojha.

## FOOTNOTES

### CONFLICTS OF INTEREST

There are no conflicts of interest to disclosure.

### FUNDING

This work was supported by Florida Department of Health (http://www.floridahealth.gov/) award number 7ZK09 and the National Institutes of Health (https://www.nih.gov/) grant number DA036154.

### CORRESPONDING AUTHOR INFORMATION

Dr. Nazira El-Hage, Ph.D., Department of Immunology and Nanomedicine, Florida International University, Herbert Wertheim College of Medicine, Miami, FL 33199, USA; E-mail address; Phone: (305)-348-4346; FAX: (305)-348-1109.

## REFERENCES

1. de Araujo TVB, Rodrigues LC, de Alencar Ximenes RA, de Barros Miranda-Filho D, Montarroyos UR, de Melo APL, Valongueiro S, de Albuquerque M, Souza WV, Braga C, Filho SPB, Cordeiro MT, Vazquez E, Di Cavalcanti Souza Cruz D, Henriques CMP, Bezerra LCA, da Silva Castanha PM, Dhalia R, Marques-Junior ETA, Martelli CMT, investigators from the Microcephaly Epidemic Research G, Brazilian Ministry of H, Pan American Health O, Instituto de Medicina Integral Professor Fernando F, State Health Department of P. 2016. Association between Zika virus infection and microcephaly in Brazil, January to May, 2016: preliminary report of a case-control study. Lancet Infect Dis 16:1356–1363.

2. Jaenisch T, Rosenberger KD, Brito C, Brady O, Brasil P, Marques ET. 2017. Risk of microcephaly after Zika virus infection in Brazil, 2015 to 2016. Bull World Health Organ 95:191–198.

3. Parra B, Lizarazo J, Jimenez-Arango JA, Zea-Vera AF, Gonzalez-Manrique G, Vargas J, Angarita JA, Zuniga G, Lopez-Gonzalez R, Beltran CL, Rizcala KH, Morales MT, Pacheco O, Ospina ML, Kumar A, Cornblath DR, Munoz LS, Osorio L, Barreras P, Pardo CA. 2016. Guillain-Barre Syndrome Associated with Zika Virus Infection in Colombia. N Engl J Med 375:1513–1523.

4. Mlakar J, Korva M, Tul N, Popovic M, Poljsak-Prijatelj M, Mraz J, Kolenc M, Resman Rus K, Vesnaver Vipotnik T, Fabjan Vodusek V, Vizjak A, Pizem J, Petrovec M, Avsic Zupanc T. 2016. Zika Virus Associated with Microcephaly. N Engl J Med 374:951–8.

5. Rasmussen SA, Jamieson DJ, Honein MA, Petersen LR. 2016. Zika Virus and Birth Defects--Reviewing the Evidence for Causality. N Engl J Med 374:1981–7.

6. Dick GW, Kitchen SF, Haddow AJ. 1952. Zika virus. I. Isolations and serological specificity. Trans R Soc Trop Med Hyg 46:509–20.

7. Barbelanne M, Tsang WY. 2014. Molecular and cellular basis of autosomal recessive primary microcephaly. Biomed Res Int 2014:547986.

8. Li C, Xu D, Ye Q, Hong S, Jiang Y, Liu X, Zhang N, Shi L, Qin CF, Xu Z. 2016. Zika Virus Disrupts Neural Progenitor Development and Leads to Microcephaly in Mice. Cell Stem Cell 19:120–6.

9. Tang H, Hammack C, Ogden SC, Wen Z, Qian X, Li Y, Yao B, Shin J, Zhang F, Lee EM, Christian KM, Didier RA, Jin P, Song H, Ming GL. 2016. Zika Virus Infects Human Cortical Neural Progenitors and Attenuates Their Growth. Cell Stem Cell 18:587–90.

10. Zhang F, Hammack C, Ogden SC, Cheng Y, Lee EM, Wen Z, Qian X, Nguyen HN, Li Y, Yao B, Xu M, Xu T, Chen L, Wang Z, Feng H, Huang WK, Yoon KJ, Shan C, Huang L, Qin Z, Christian KM, Shi PY, Xu M, Xia M, Zheng W, Wu H, Song H, Tang H, Ming GL, Jin P. 2016. Molecular signatures associated with ZIKV exposure in human cortical neural progenitors. Nucleic Acids Res 44:8610–8620.

11. Faheem M, Naseer MI, Rasool M, Chaudhary AG, Kumosani TA, Ilyas AM, Pushparaj P, Ahmed F, Algahtani HA, Al-Qahtani MH, Saleh Jamal H. 2015. Molecular genetics of human primary microcephaly: an overview. BMC Med Genomics 8 Suppl 1:S4.

12. Gilmore EC, Walsh CA. 2013. Genetic causes of microcephaly and lessons for neuronal development. Wiley Interdiscip Rev Dev Biol 2:461–78.

13. Ojha CR, Rodriguez M, Karuppan MKM, Lapierre J, Kashanchi F, El-Hage N. 2019. Toll-like receptor 3 regulates Zika virus infection and associated host inflammatory response in primary human astrocytes. PLoS One 14:e0208543.

14. Klionsky DJ, Baehrecke EH, Brumell JH, Chu CT, Codogno P, Cuervo AM, Debnath J, Deretic V, Elazar Z, Eskelinen EL, Finkbeiner S, Fueyo-Margareto J, Gewirtz D, Jaattela M, Kroemer G, Levine B, Melia TJ, Mizushima N, Rubinsztein DC, Simonsen A, Thorburn A, Thumm M, Tooze SA. 2011. A comprehensive glossary of autophagy-related molecules and processes (2nd edition). Autophagy 7:1273–94.

15. Lapierre J, Rodriguez M, Ojha CR, El-Hage N. 2018. Critical Role of Beclin1 in HIV Tat and Morphine-Induced Inflammation and Calcium Release in Glial Cells from Autophagy Deficient Mouse. J Neuroimmune Pharmacol 13:355–370.

16. Rodriguez M, Kaushik A, Lapierre J, Dever SM, El-Hage N, Nair M. 2017. Electro-Magnetic Nano-Particle Bound Beclin1 siRNA Crosses the Blood-Brain Barrier to Attenuate the Inflammatory Effects of HIV-1 Infection in Vitro. J Neuroimmune Pharmacol 12:120–132.

17. Rodriguez M, Lapierre J, Ojha CR, Kaushik A, Batrakova E, Kashanchi F, Dever SM, Nair M, El-Hage N. 2018. Author Correction: Intranasal drug delivery of small interfering RNA targeting Beclin1 encapsulated with polyethylenimine (PEI) in mouse brain to achieve HIV attenuation. Scientific reports 8:4778–4778.

18. Gurwell JA, Nath A, Sun Q, Zhang J, Martin KM, Chen Y, Hauser KF. 2001. Synergistic neurotoxicity of opioids and human immunodeficiency virus-1 Tat protein in striatal neurons in vitro. Neuroscience 102:555–63.

19. Macdonald J, Tonry J, Hall RA, Williams B, Palacios G, Ashok MS, Jabado O, Clark D, Tesh RB, Briese T, Lipkin WI. 2005. NS1 protein secretion during the acute phase of West Nile virus infection. J Virol 79:13924–33.

20. Yu C, Wang L, Lv B, Lu Y, Zeng Le, Chen Y, Ma D, Shi T, Wang L. 2008. TMEM74, a lysosome and autophagosome protein, regulates autophagy. Biochemical and Biophysical Research Communications 369:622–629.

21. Tripathi S, Balasubramaniam VR, Brown JA, Mena I, Grant A, Bardina SV, Maringer K, Schwarz MC, Maestre AM, Sourisseau M, Albrecht RA, Krammer F, Evans MJ, Fernandez-Sesma A, Lim JK, Garcia-Sastre A. 2017. A novel Zika virus mouse model reveals strain specific differences in virus pathogenesis and host inflammatory immune responses. PLoS Pathog 13:e1006258.

22. Lazear HM, Govero J, Smith AM, Platt DJ, Fernandez E, Miner JJ, Diamond MS. 2016. A Mouse Model of Zika Virus Pathogenesis. Cell Host Microbe 19:720–30.

23. Wu KY, Zuo GL, Li XF, Ye Q, Deng YQ, Huang XY, Cao WC, Qin CF, Luo ZG. 2016. Vertical transmission of Zika virus targeting the radial glial cells affects cortex development of offspring mice. Cell Res 26:645–54.

24. Schweighardt B, Atwood WJ. 2001. Glial cells as targets of viral infection in the human central nervous system, p 721–735, Progress in Brain Research, vol 132. Elsevier.

25. Furr SR, Marriott I. 2012. Viral CNS infections: role of glial pattern recognition receptors in neuroinflammation. Frontiers in Microbiology 3:201.

26. Anfasa F, Siegers JY, van der Kroeg M, Mumtaz N, Stalin Raj V, de Vrij FMS, Widagdo W, Gabriel G, Salinas S, Simonin Y, Reusken C, Kushner SA, Koopmans MPG, Haagmans B, Martina BEE, van Riel D. 2017. Phenotypic Differences between Asian and African Lineage Zika Viruses in Human Neural Progenitor Cells. mSphere 2.

27. Alcon S, Talarmin A, Debruyne M, Falconar A, Deubel V, Flamand M. 2002. Enzyme-linked immunosorbent assay specific to Dengue virus type 1 nonstructural protein NS1 reveals circulation of the antigen in the blood during the acute phase of disease in patients experiencing primary or secondary infections. J Clin Microbiol 40:376–81.

28. Libraty DH, Young PR, Pickering D, Endy TP, Kalayanarooj S, Green S, Vaughn DW, Nisalak A, Ennis FA, Rothman AL. 2002. High circulating levels of the dengue virus nonstructural protein NS1 early in dengue illness correlate with the development of dengue hemorrhagic fever. J Infect Dis 186:1165–8.

29. Cao B, Parnell LA, Diamond MS, Mysorekar IU. 2017. Inhibition of autophagy limits vertical transmission of Zika virus in pregnant mice. J Exp Med 214:2303–2313.

30. Ferdin J, Goricar K, Dolzan V, Plemenitas A, Martin JN, Peterlin BM, Deeks SG, Lenassi M. 2018. Viral protein Nef is detected in plasma of half of HIV-infected adults with undetectable plasma HIV RNA. PLoS One 13:e0191613.

31. Merfeld E, Ben-Avi L, Kennon M, Cerveny KL. 2017. Potential mechanisms of Zika-linked microcephaly. Wiley Interdiscip Rev Dev Biol 6.

32. Nicholas AK, Khurshid M, Désir J, Carvalho OP, Cox JJ, Thornton G, Kausar R, Ansar M, Ahmad W, Verloes A, Passemard S, Misson J-P, Lindsay S, Gergely F, Dobyns WB, Roberts E, Abramowicz M, Woods CG. 2010. WDR62 is associated with the spindle pole and is mutated in human microcephaly. Nature genetics 42:1010–1014.

33. Yu TW, Mochida GH, Tischfield DJ, Sgaier SK, Flores-Sarnat L, Sergi CM, Topçu M, McDonald MT, Barry BJ, Felie J, Sunu C, Dobyns WB, Folkerth RD, Barkovich AJ, Walsh CA. 2010. Mutations in WDR62, encoding a centrosome-associated protein, cause microcephaly with simplified gyri and abnormal cortical architecture. Nature genetics 42:1015–1020.

34. Chen J-F, Zhang Y, Wilde J, Hansen KC, Lai F, Niswander L. 2014. Microcephaly disease gene Wdr62 regulates mitotic progression of embryonic neural stem cells and brain size. Nature Communications 5:3885.

35. Szczepanski S, Hussain MS, Sur I, Altmüller J, Thiele H, Abdullah U, Waseem SS, Moawia A, Nürnberg G, Noegel AA, Baig SM, Nürnberg P. 2016. A novel homozygous splicing mutation of CASC5 causes primary microcephaly in a large Pakistani family. Human Genetics 135:157–170.

36. Mathew R, Kongara S, Beaudoin B, Karp CM, Bray K, Degenhardt K, Chen G, Jin S, White E. 2007. Autophagy suppresses tumor progression by limiting chromosomal instability. Genes & Development 21:1367–1381.

37. Zhao Z, Oh S, Li D, Ni D, Pirooz Sara D, Lee J-H, Yang S, Lee J-Y, Ghozalli I, Costanzo V, Stark Jeremy M, Liang C. 2012. A Dual Role for UVRAG in Maintaining Chromosomal Stability Independent of Autophagy. Developmental Cell 22:1001–1016.

38. Kadir R, Harel T, Markus B, Perez Y, Bakhrat A, Cohen I, Volodarsky M, Feintsein-Linial M, Chervinski E, Zlotogora J, Sivan S, Birnbaum RY, Abdu U, Shalev S, Birk OS. 2016. ALFY-Controlled DVL3 Autophagy Regulates Wnt Signaling, Determining Human Brain Size. PLOS Genetics 12:e1005919.

39. Boland B, Kumar A, Lee S, Platt FM, Wegiel J, Yu WH, Nixon RA. 2008. Autophagy induction and autophagosome clearance in neurons: relationship to autophagic pathology in Alzheimer’s disease. J Neurosci 28:6926–37.

40. Chiramel AI, Best SM. 2018. Role of autophagy in Zika virus infection and pathogenesis. Virus Res 254:34–40.

41. Rodriguez M, Lapierre J, Ojha CR, Estrada-Bueno H, Dever SM, Gewirtz DA, Kashanchi F, El-Hage N. 2017. Importance of Autophagy in Mediating Human Immunodeficiency Virus (HIV) and Morphine-Induced Metabolic Dysfunction and Inflammation in Human Astrocytes. Viruses 9.

42. Houtman J, Freitag K, Gimber N, Schmoranzer J, Heppner FL, Jendrach M. 2019. Beclin1-driven autophagy modulates the inflammatory response of microglia via NLRP3. EMBO J 38.

43. Dever SM, Rodriguez M, Lapierre J, Costin BN, El-Hage N. 2015. Differing roles of autophagy in HIV-associated neurocognitive impairment and encephalitis with implications for morphine co-exposure. Front Microbiol 6:653.

44. Cao B, Parnell LA, Diamond MS, Mysorekar IU. 2017. Inhibition of autophagy limits vertical transmission of Zika virus in pregnant mice. The Journal of Experimental Medicine 214:2303–2313.

45. Ranke MB, Wolfle J, Schnabel D, Bettendorf M. 2009. Treatment of dwarfism with recombinant human insulin-like growth factor-1. Dtsch Arztebl Int 106:703–9.

46. Giabicani E, Chantot-Bastaraud S, Bonnard A, Rachid M, Whalen S, Netchine I, Brioude F. 2019. Roles of Type 1 Insulin-Like Growth Factor (IGF) Receptor and IGF-II in Growth Regulation: Evidence From a Patient Carrying Both an 11p Paternal Duplication and 15q Deletion. Front Endocrinol (Lausanne) 10:263.

47. Beck KD, Powell-Braxton L, Widmer HR, Valverde J, Hefti F. 1995. Igf1 gene disruption results in reduced brain size, CNS hypomyelination, and loss of hippocampal granule and striatal parvalbumin-containing neurons. Neuron 14:717–30.

48. Riikonen R, Makkonen I, Vanhala R, Turpeinen U, Kuikka J, Kokki H. 2006. Cerebrospinal fluid insulin-like growth factors IGF-1 and IGF-2 in infantile autism. Dev Med Child Neurol 48:751–5.

49. Riikonen R. 2006. Insulin-like growth factor delivery across the blood-brain barrier. Potential use of IGF-1 as a drug in child neurology. Chemotherapy 52:279–81.

50. Riikonen R. 2017. Insulin-Like Growth Factors in the Pathogenesis of Neurological Diseases in Children. Int J Mol Sci 18.

51. Badadani M. 2012. Autophagy Mechanism, Regulation, Functions, and Disorders %J ISRN Cell Biology. 2012:11.

52. Zhao Z, Yang M, Azar SR, Soong L, Weaver SC, Sun J, Chen Y, Rossi SL, Cai J. 2017. Viral Retinopathy in Experimental Models of Zika Infection. Invest Ophthalmol Vis Sci 58:4355–4365.

53. Toborek M, Lee YW, Pu H, Malecki A, Flora G, Garrido R, Hennig B, Bauer HC, Nath A. 2003. HIV-Tat protein induces oxidative and inflammatory pathways in brain endothelium. J Neurochem 84:169–79.

54. Glass CK, Saijo K, Winner B, Marchetto MC, Gage FH. 2010. Mechanisms underlying inflammation in neurodegeneration. Cell 140:918–34.

55. Jha MK, Jeon S, Suk K. 2012. Glia as a Link between Neuroinflammation and Neuropathic Pain. Immune Netw 12:41–7.

